# NetGAM: Using generalized additive models to improve the predictive power of ecological network analyses constructed using time-series data

**DOI:** 10.1101/2021.10.22.465515

**Authors:** Samantha J. Gleich, Jacob A. Cram, Jake L. Weissman, David A. Caron

## Abstract

Ecological network analyses are used to identify potential biotic interactions between microorganisms from species abundance data. These analyses are often carried out using time-series data; however, time-series networks have unique statistical challenges. Time-dependent species abundance data can lead to species co-occurrence patterns that are not a result of direct, biotic associations and may therefore result in inaccurate network predictions. Here, we describe a generalize additive model (GAM)-based data transformation that removes time-series signals from species abundance data prior to running network analyses. Validation of the transformation was carried out by generating mock, time-series datasets, with an underlying covariance structure, running network analyses on these datasets with and without our GAM transformation, and comparing the network outputs to the known covariance structure of the simulated data. The results revealed that seasonal abundance patterns substantially decreased the accuracy of the inferred networks. Additionally, the GAM transformation increased the F1 score of inferred ecological networks on average and improved the ability of network inference methods to capture important features of network structure. This study underscores the importance of considering temporal features when carrying out network analyses and describes a simple, effective tool that can be used to improve results.

## Introduction

Communities of microorganisms exist in virtually all natural environments on the planet and are shaped by complex interactions. Species-species interactions are diverse and may benefit both species involved (e.g. mutualism), only one (e.g. commensalism, parasitism), or hurt both (e.g. competition)[1–2]. Ecological network analyses are increasingly used by microbial ecologists to identify potential biotic interactions between organisms and to form hypotheses regarding microbial community structure and function [1, 3–4]. Commonly employed network-inference methods include pairwise correlation-based, regression-based, and probabilistic graphical methods [5–6]. All of these methods leverage microbial abundance measurements to identify co-occurrence patterns between organisms [1, 7–8]. Biotic interactions between organisms are then predicted based on the species’ co-occurrence patterns, resulting in a list of nodes (organisms) connected by edges (associations) [1, 7–9].

Abundance data that are used as input for ecological network analyses are often obtained through monthly time-series sampling efforts [10–15]. Time-series datasets are valuable because documenting changes in microbial community structure over long timeframes can provide information on the monthly, annual, or interannual variability in species abundances [16–17]. For example, ~11 years of monthly sampling at the San Pedro Ocean Time-Series (SPOT) site revealed that 23% of bacterial operational taxonomic units (OTUs) demonstrated predictable, seasonal abundance patterns in surface waters [13]. Another time-series study that took place over ~6 years in the Western English Channel revealed that the month of year could explain over half of the variability in bacterial community composition [12]. Such studies highlight how long-term time-series datasets may be used to identify predictable changes in microbial community composition over time.

Time-series datasets can provide information on microbial community composition and structure, but ecological networks inferred from them should be built and interpreted with caution. There are statistical challenges associated with time-series network analyses because the samples are not independent over time [12, 18–19]. This inherent time-dependence may be influenced in part by seasonally reoccurring patterns in the abiotic environment (e.g. seasonal mixing or upwelling events) or long-term changes in the environment over-time (e.g. rising seawater temperature) [17, 20–21]. As a result of this time-dependence, species abundance patterns can lead to co-occurrence patterns that yield spurious network predictions [4, 22]. For instance, two species may both attain maximal abundances during spring nutrient upwelling events, even if no interactions occur between them. Their shared periodicity in this case may manifest itself as a ‘false association’. Time-dependence may also confound species co-occurrence patterns through the effects of Simpson’s paradox [23–24]. If a mutualistic relationship exists between two species, one might expect a positive correlation between the abundances of the two species across samples. However, if the species respond differently to a third variable (e.g. month of year), then the positive association between the two species may be offset or reversed as a result of this time-dependence [24]. Such inaccurate associations indicate that caution should be exercised when carrying out network analyses on time-series datasets [7, 24].

Here, we propose and validate a generalized additive model (GAM)-based data transformation that corrects for potentially confounding time-series signals that are prevalent in microbial relative abundance data. The GAM transformation is conducted prior to carrying out ecological network analyses, and removes seasonal, long-term, and autocorrelative trends, thereby allowing researchers to focus on the residual statistical variability of the microbial abundance data. We contend that the residual variability is likely more indicative of true biotic associations than are untransformed data. We used GAMs in this data transformation method, as they are versatile, and commonly used to capture non-linear trends typical of time-series data [25]. Generalized additive models have been used to model both seasonal patterns and long-term trends in time-series data [25–27] and have also been used to capture autocorrelative signals [17, 28]. The GAM-based data transformation presented here has the potential to capture seasonal, long-term, and autocorrelative trends in time-series datasets, thus minimizing the influence that temporal signals have on inferred microbial co-occurrence patterns and increasing the predictive power of commonly employed networking methods.

## Materials and Methods

Our general strategy was to compare the performance of 4 approaches for inferring microbial associations from abundance data with overlying time-series signals. The approaches were (1) pairwise spearman correlation analysis (SCC) [1, 29], (2) Graphical lasso analysis (Glasso) [30–31], (3) pairwise SCC analysis with a pre-processing step where seasonal and long-term splines were fit to and subtracted from each variable using a GAM (GAM-SCC), and (4) Glasso with the same GAM subtraction approach (GAM-Glasso). Our validation strategy for the GAM transformation consisted of generating mock datasets with underlying associations, masking those associations by adding seasonal and long-term signals to the abundance data, and comparing the predicted associations obtained from each network inference method to the true species-species associations.

### Data simulation: Generating mock abundance data with time-series properties

We generated mock abundance datasets that had a predetermined, underlying network structure and contained long-term and seasonal species abundance patterns. First, a covariance matrix was generated to describe the relationships between species in a mock network (Figure S1, Panel 1). The covariance matrices were constructed with underlying network structures that followed either a Barabási-Albert model, which have scale-free properties, an Erdős-Rényi model, in which connections between nodes are random, or a model of network topology based on a real microbial dataset (American Gut Project dataset; Figure S1) [32–33]. The Erdős-Rényi and Barabási-Albert model datasets were generated so that each dataset contained 400 species and 200 samples, and the American Gut model datasets were created so that each dataset contained 127 species and 200 samples. A random Bernoulli distribution was used to simulate the covariance matrix for the Erdős-Rényi networks. We set the probability of interactions occurring between species in a given Erdős-Rényi network to 1%, making the simulated networks 99% sparse. The Barabási-Albert networks were generated using the “sample_pa” function in the igraph package [34]. The “graph2prec” function in the SpiecEasi package was used to predict the covariance matrix of the American Gut Project dataset [33]. The covariance between species in a dataset was considered “high” when true associations in the covariance matrix were set to 100. Conversely, the covariance between species was considered “low” if the true associations in the covariance matrix were set to 10. These covariance matrices describe the “real”, underlying species interactions in our mock species abundance datasets.

After generating a covariance matrix using the Barabási-Albert, Erdős-Rényi, or American Gut Project model, the mean abundance for each species was generated from a normal distribution with a mean of 10 and a variance of 1. These mean abundance values and the covariance matrix were used to parameterize a multivariate normal distribution from which species abundance values for all 200 samples in a dataset were drawn (Figure S1, Panel 2). The values generated from this multivariate normal distribution were the species abundance values without time-series features confounding the relationship between 2 associated species.

Seasonal trends were added to 0%, 25%, 50% or 100% of the species in the mock networks. The seasonal signals were generated by plugging a vector of consecutive integers of length 200 (*N*_*t*_) into the “gradual” (Eqn 1.) or “abrupt” (Eqn 2.) seasonal equations (Figure S1, Panel 3). The starting value of this vector of consecutive integers was drawn at random to allow seasonal peaks centered at different months for different species…

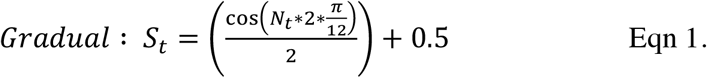

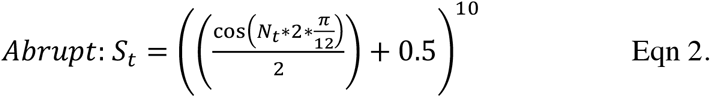

…where *N* is the random vector of consecutive integers, *S* is the output seasonal vector, and *t* is the index of vectors *N* and *S.* Each element in the seasonal vector (*S_t_*) was then multiplied by the corresponding element in the abundance vector (*X_t_*) of a specific species to obtain mock species abundance values with a “gradual” or “abrupt” seasonal trend.

A long-term time-series trend was added to the abundance values of 0% or 50% of the species in the mock datasets (Figure S1, Panel 4). When a long-term signal was applied to 50% of the species in a dataset, half of the species were randomly selected to have this long-term trend. Then, a vector of linear values was generated following Eqn 3, such that

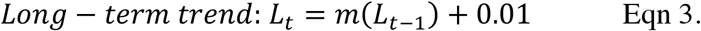

*L_t_* is the point in the line at the next time point and *m* is the slope of the line. The slope parameter (*m*) was generated from a random normal distribution with a mean of 0.01 and a variance of 0.01. Half of the long-term trends generated had a positive slope (increasing over time) and half had a negative slope (decreasing over time). After generating the vector of linear values (*L_t_*), each element of this vector was added to each element of the abundance vector (*X_t_*) of a specific species to simulate long-term time-series trends for selected species.

Time-series predictor columns were added to each dataset after applying monthly and long-term abundance trends to a portion of the species in the mock datasets. The predictor columns that were used in the downstream GAM-based data transformation were the month of the year (i.e. 1-12) and the day of the time-series (i.e. 1-200). In total, we generated 100 mock datasets for every combination of conditions (Table S1), resulting in 8 400 mock time-series datasets that were used in the downstream GAM transformation and network analysis procedures.

### Data simulation: Simulating count data from abundance values

The time-series species abundance data were transformed to make the abundance values resemble high-throughput sequencing data because microbial time-series sampling efforts are often processed using such molecular methods (e.g. tag-sequencing, meta-omics). Relative abundances of different species in natural communities are highly skewed, so that relatively few species constitute most of the organisms in a sample although many rare species are also present [35–36]. To transform abundance data into sequence data that look like a natural community, species abundances were first exponentiated to increase the prevalence of abundant species and to decrease the prevalence of rare species (Figure S1, Panel 5). These exponentiated species abundance values were then converted to relative abundance values because microbial abundances are dependent on sequencing depth in high-throughput sequencing studies and are therefore inherently relative (i.e. compositional) [37]. The relative abundance values were calculated by dividing each species count by the sum of all species counts in each sample (Figure S1, Panel 6). The resulting relative abundance values and time-series predictor variables were then used in data normalization and GAM-transformation steps prior to carrying out the network analyses.

### Network inference: Count data normalization and GAM transformation

A variety of steps were taken to reverse the above transformation steps and back out the species-species relationships in the underlying networks. We advocate these steps to infer network structure from a real time-series dataset. A centered log-ratio (CLR) transformation was first applied to the species relative abundance values to normalize the mock species abundance data across samples using the “clr” function in the compositions package in R (Figure 1) [38]. This transformation step is important to avoid spurious inferences induced by the inherent compositionality of relative abundance data [31, 33, 37]. The CLR-transformed dataset was copied, with one copy subjected to a subsequent GAM transformation, and the other one not GAM-transformed.

**Figure 1:**
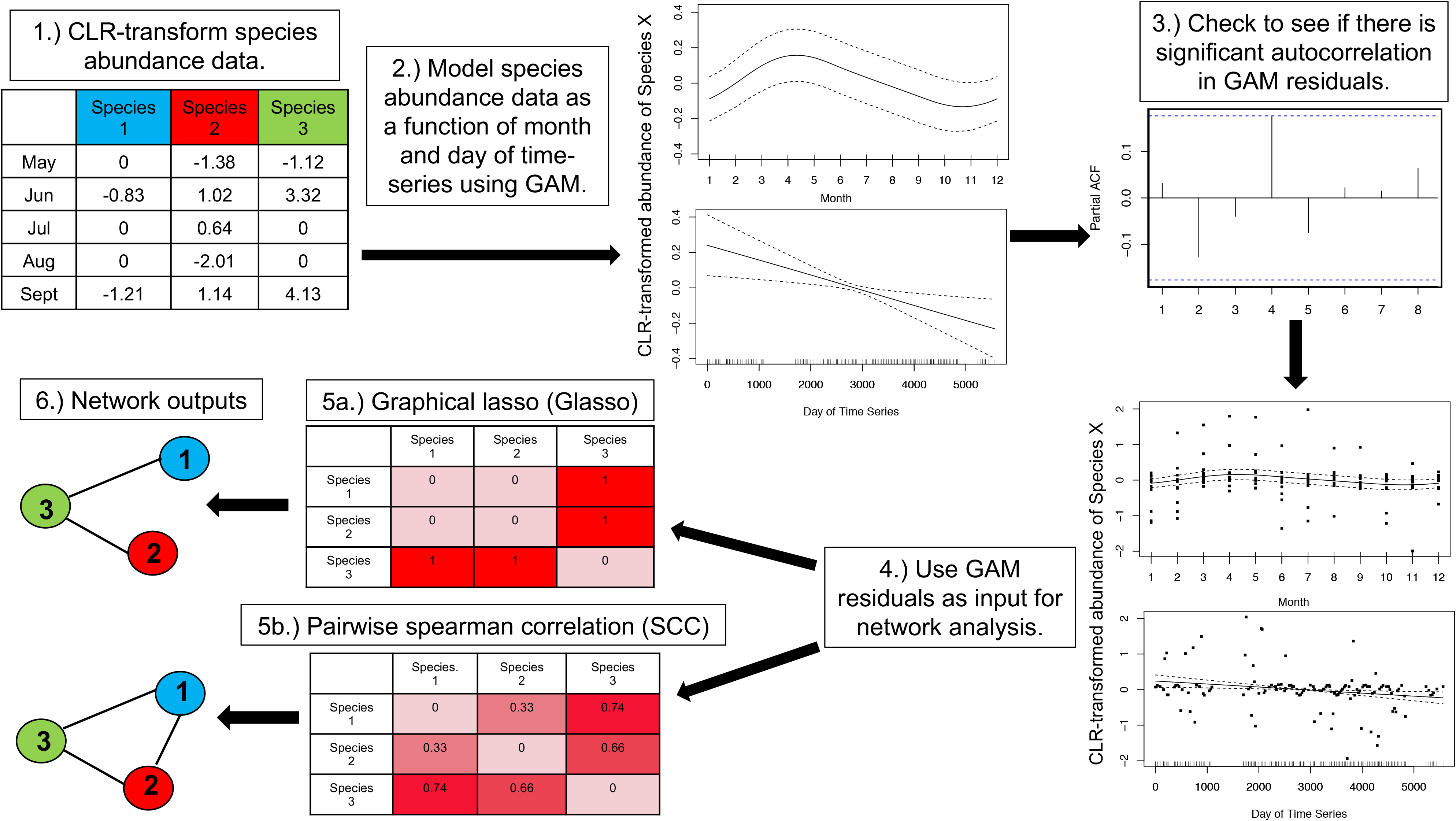
Steps used to carry out the GAM-based transformation of time-series species abundance data prior to carrying out pairwise spearman correlation (SCC) and graphical lasso (Glasso) ecological network analyses. The raw, species abundance data were first CLR-transformed (1). Generalized additive models (GAMs) were then fit to each species in the dataset (2) and the residuals of each GAM were checked for significant autocorrelation (3). The residuals of each GAM were extracted (4) and were used as input in the SCC and Glasso network analysis methods (5). Finally, the GAM-transformed network outputs were obtained (6; see text for additional details).

The GAM-transformation was carried out by fitting GAMs to each individual species in the dataset to remove monthly signals, long-term trends, and autocorrelation from the species abundance data. These GAMs were fit using the “gamm” function in the mgcv package in R [39–40]. The GAMs that were used included the “month of year” parameter as a cyclical spline predictor and the “day of time-series” parameter as a thin-plate spline predictor (Figure 1). Additionally, the first GAM included a continuous AR1 correlation structure term in the model. This GAM was revised for specific species in the dataset when the GAM could not be resolved or when significant autocorrelation was detected in the GAM residuals (Figure 1). This GAM revision step tested a number of different correlation structure terms (i.e. AR1, CompSymm, Exp, and Gaus) in the models and is automated in the NetGAM package, which we have published. After fitting a GAM to all of the species in the input dataset, the residuals of each GAM were extracted and were used as the new, GAM-transformed abundance values for each species (Figure 1). These GAM residuals represent species abundance values with a reduced influence of time and were used as input in the downstream GAM-SCC and GAM-Glasso network analyses.

### Network inference: Network runs and statistical analyses

The pre-processed species abundance data with and without the GAM-removal of time-series signals were used in the 2 networking methods of interest (SCC and Glasso) in order to compare the outputs of the 4 network inference approaches (SCC, GAM-SCC, Glasso, and GAM-Glasso). A nonparanormal transformation was first applied to the species abundance datasets with and without GAM transformation using the “huge.npn” function in the huge package in R [41]. Spearman correlation networks (SCC, GAM-SCC) were then constructed by calculating the correlation between every pair of species in the mock abundance datasets. A Bonferroni-corrected *p*-value of 0.01 was used as a cutoff to identify edges in these SCC networks. The Glasso networks (Glasso, GAM-Glasso) were constructed by testing 30 regularization parameter values (i.e. lambdas) in each network using the “batch.pulsar” (criterion = “stars”; rep.num = 50) function in the pulsar package in R [42]. The lambda that resulted in the most stable network output was selected using the StARS method [43]. Finally, the graph that resulted from the StARS output was used to obtain a species adjacency matrix for the Glasso networks with and without GAM transformation.

The Glasso adjacency matrices and the SCC results were used to generate lists of species-species associations predicted by the network outputs. The species-species associations predicted by the networks were then compared to the true species-species associations and the F1 scores of the network predictions were calculated. The F1 score is a measure of classification performance (presence or absence of an edge) that, unlike accuracy, accounts for uneven classes, which is essential when dealing with sparse networks. The F1 scores of the GAM-transformed networks (GAM-SCC, GAM-Glasso) were compared to the networks that did not undergo GAM transformation (SCC, Glasso) using paired Wilcoxon tests with Bonferroni correction. An adjusted *p*-value of 0.01 was used as a cutoff to identify under what circumstances the GAM significantly improved the F1 score of a Glasso or SCC network. Information about the effectiveness of the GAM transformation in improving the predictive power of the time-series networks was thereby obtained.

### Network inference: Comparison of predicted network structures

Additional networks were generated using the methods described above to compare the predicted network structures obtained from the four network approaches (GAM-Glasso, Glasso, GAM-SCC, and SCC) to the real network structures. The average clustering coefficient and the degree distribution of these additional network outputs were calculated and used for the network structure comparisons. The average clustering coefficient of a network describes the likelihood that two species that are both associated with a third species are also associated with each other [44], and in a sense describes the “clumpiness” of a network. The network degree distributions describe the probability distribution of the number of interactions per node in a network [45].

The network structure comparisons were carried out by generating 100 additional Barabási-Albert and Erdős-Rényi datasets with high species-species covariance. Each of the mock, time-series datasets that were used as input in these additional network runs contained 100 species and 200 samples. Additionally, 50% of the species in each of these datasets had a gradual, seasonal abundance pattern (Figure S1, Panel 3). The SCC, GAM-SCC, Glasso, and GAM-Glasso networking approaches were used to obtain network predictions for these extra network iterations and the average precision, recall, and F1 score obtained from each of the four approaches was calculated. The average clustering coefficients of the networks were also calculated and were compared to the average clustering coefficients of the real Barabási-Albert and Erdős-Rényi networks. Finally, the degree distributions of the network runs were calculated and plotted alongside to the real network degree distributions.

## Results

### Seasonal abundance patterns decreased the performance of network inference methods

The F1 scores of the reconstructed network outputs generally decreased (indicating worse performance) as the proportion of species in the mock dataset with a seasonal abundance pattern increased. The decreases in network F1 scores with increases in the percent of species with a seasonal abundance pattern were always prevalent in the Glasso and SCC networks when the GAM transformation was not applied and when some percentage of species (> 0%) in the mock datasets had a seasonal abundance pattern (Figure 2, Tables S2-S4). In general, the highest F1 scores were associated with networks that did not contain any species with an underlying seasonal signal (0%), while the lowest F1 scores were typically associated with networks in which all of the species had a seasonal abundance pattern (100%; Figure 2, Tables S2-S4).

**Figure 2:**
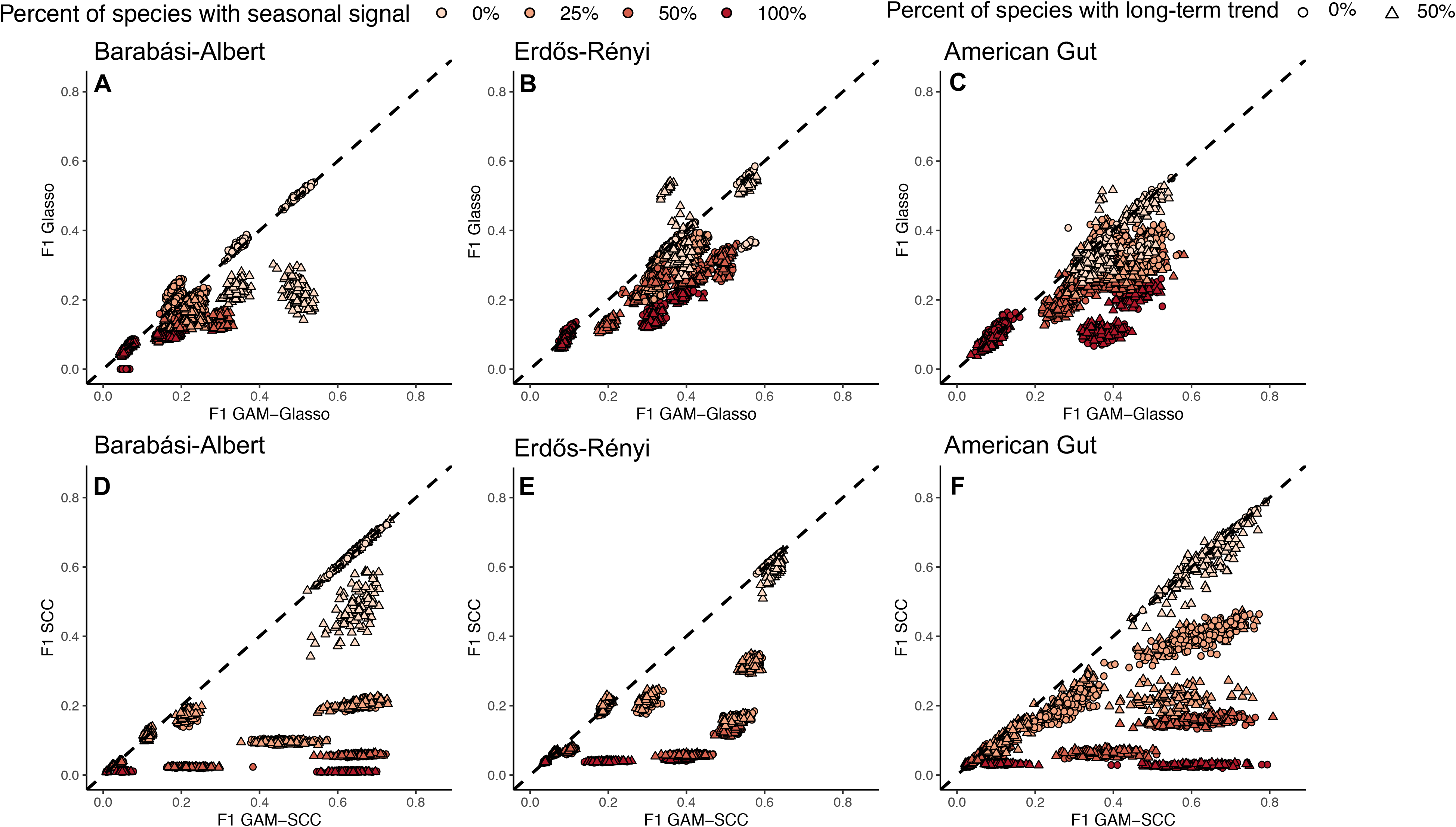
F1 scores of the networks constructed without GAM transformation (y-axis) plotted against the F1 scores of the GAM-transformed networks (x-axis) for all of the mock time-series datasets that were simulated. Panels A-C show the comparison between the Glasso and the GAM-Glasso networks, while panels D-F show the comparison between the SCC and the GAM-SCC networks. The dashed, black lines show the 1:1 relationship. Data points below the 1:1 line depict network outputs that had a higher F1 score after applying the GAM-based data transformation, while data points that fall above the 1:1 line depict network runs that had a higher F1 score without applying the GAM-based data transformation prior to network construction.

The general decline in network F1 score with a greater percentage of species exhibiting seasonality was often less pronounced when the mock datasets were GAM-transformed prior to carrying out the network analyses. For example, when species-species covariance was high, the GAM-SCC method tended to perform similarly regardless of whether 25%, 50%, or 100% of the species in a Barabási-Albert or American Gut dataset network had a gradual seasonal abundance pattern (Figure S3 and S7; Panels B and F). The decline in network F1 score with increases in the percentage of species exhibiting seasonality was also less pronounced in GAM-Glasso networks when there was high covariance between species and when some of the species in the input dataset had a gradual, seasonal abundance pattern (Figures S2, S4, and S6; Panels B and F).

### GAM transformation improved network inference on average

The GAM transformation increased the F1 score of the Glasso networks in 82.0% of all network runs (Figure 2; Panels A-C; most points fall below the 1:1 line). Specifically, the GAM transformation significantly increased the mean F1 score of the Glasso networks when a gradual seasonal signal (Figure S1; Panel 3) was applied to some fraction of the species in the input dataset (Figures S2, S4, and S6; Panels B, D, F, and H). The GAM-Glasso networks also had significantly higher F1 scores when 50% of species in the input dataset had an abrupt, seasonal abundance pattern (Figures S2, S4, and S6; Panels A, C, E, and G). For the Barabási-Albert models, the F1 scores of the GAM-Glasso networks were significantly greater than those of the Glasso networks when 50% of the species in the input dataset had a long-term trend with no (0%) seasonality (Figure S2; Panels E-H). Similar increases in F1 score were noted in the Erdős-Rényi and American Gut dataset, GAM-Glasso networks when the covariance between species was low and when 50% of the species in the dataset had long-term changes in abundance with no (0%) seasonality (Figure S4 and S6; Panels G and H).

The GAM transformation also led to substantial increases in the F1 score of the SCC networks. Overall, the GAM transformation increased the F1 score of SCC networks in 79.3% of all network runs (Figure 2; Panels D-F; most points fall below the 1:1 line). The average F1 scores of the networks were significantly greater when the data were GAM-transformed prior to carrying out a SCC network analysis when a gradual seasonal signal (Figure S1; Panel 3) was applied to some fraction of the species in the input dataset (Figures S3, S5, and S7; Panels B, D, F, and H). The mean F1 scores of all GAM-SCC networks were also significantly greater than those of the SCC networks when there was high covariance between species and when an abrupt seasonal signal (Figure S1, Panel 3) was applied to 25% or 50% of the species in the input dataset (Figures S3, S5, and S7; Panels A and E).

### GAM transformation improved the ability of Glasso and SCC networks to capture real network structure

The GAM-transformation improved the ability of the Glasso and SCC methods to capture the underlying structure of the Barabási-Albert and Erdős-Rényi networks (Figures 3–4). The real network degree distributions were more similar to the GAM-SCC and GAM-Glasso degree distributions than they were to the degree distributions of the SCC and Glasso networks without GAM transformation (Figures 3–4). The GAM-SCC approach was the most successful in capturing the real, scale-free Barabási-Albert network degree distribution and had the highest average precision, recall, and F1 score of the 4 methods tested (Figure 3; Panel C). Conversely, the GAM-Glasso approach did the best job of capturing the real, Erdős-Rényi network structure, as the SCC and GAM-SCC approaches predicted a number of high-degree nodes that were not present in the true, network structures (Figure 4; long right tail on SCC and GAM-SCC plots). Some high-degree nodes were also predicted in the Glasso and GAM-Glasso, Erdős-Rényi networks (Figure 4; long right tail on Glasso and GAM-Glasso plots) but in general were less pronounced than those noted in the SCC and GAM-SCC degree distributions.

**Figure 3:**
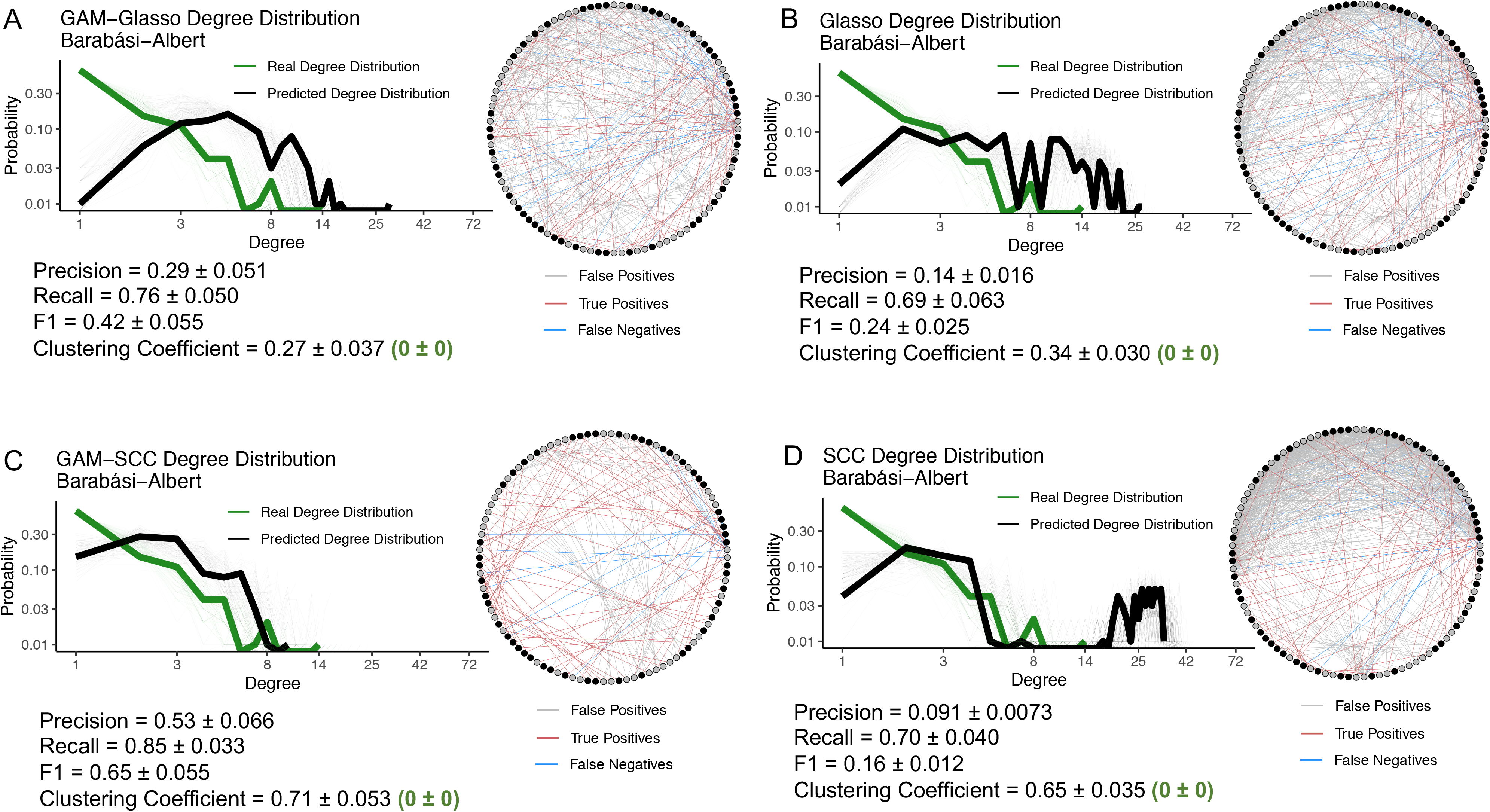
The GAM-SCC networks did the best job of capturing the real, Barabási-Albert network degree distribution. The degree distributions and network outputs of 100 GAM-Glasso (A), Glasso (B), GAM-SCC (C), and SCC (D) time-series networks constructed with 100 species. The networks depicted were constructed with mock species abundance data that had an underlying Barabási-Albert network structure and that contained 50 species with a gradual, seasonal abundance pattern. The fine green lines on the log-log plots show the real degree distributions and the fine black lines show the network-predicted degree distributions. The bolded lines show the degree distributions of the representative networks that are depicted. On the representative network images, the red edges show those edges that are true positive associations, the blue edges show those edges that are false negative associations, and the grey edges show those edges that are false positive associations. The black nodes in the network images represent the species that have a seasonal abundance pattern, while the grey nodes represent those species that do not have a seasonal abundance pattern.

**Figure 4:**
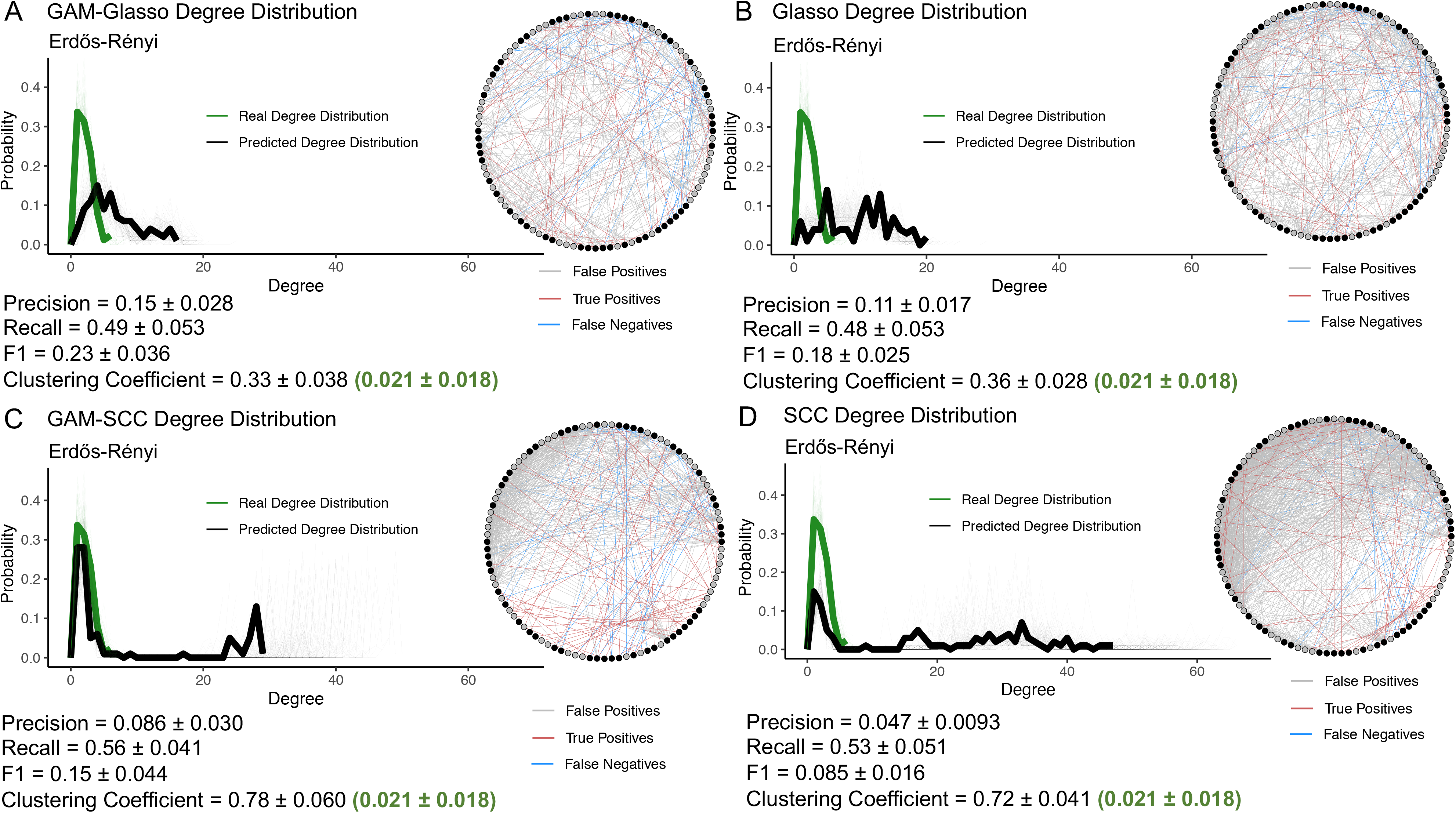
The GAM-Glasso networks did the best job of capturing the real, Erdős-Rényi network topology. The degree distributions and network outputs of 100 GAM-Glasso, Glasso, GAM-SCC, and SCC time-series networks constructed with 100 species. The networks depicted were constructed with mock species abundance data that had an underlying Erdős-Rényi network structure and that contained 50 species with a gradual, seasonal abundance pattern. Panels and color coding are the same as described for Figure 3.

The average clustering coefficient of the GAM-Glasso networks was the most similar to the average clustering coefficient of the real networks, while the average clustering coefficient of the GAM-SCC networks was the least similar to that of the real networks (Figures 3–4). The exaggerated average clustering coefficients of the SCC and GAM-SCC networks were to be expected, given the transitive nature of correlative relationships between variables. In general, the average clustering coefficient values that were obtained from the 4 network prediction approaches were substantially higher than the average clustering coefficient values of the real networks regardless of whether the underlying network structure was that of a Barabási-Albert or an Erdős-Rényi model. These high, average clustering coefficient values implied that the outputs obtained from the network inference approaches resulted in networks that were “clumpier” than the real networks.

### Effectiveness of the GAM transformation decreased when seasonal abundance patterns were abrupt

The GAM transformation slightly decreased the predictive power of a small subset of the Glasso networks when abrupt seasonal abundance patterns (Figure S1; Panel 3) were prevalent in the input dataset (Figure S2, S4, and S6; Panels A, C, E, and G). Specifically, when the covariance between species was high and all of the species (100%) in the input dataset had an abrupt seasonal abundance pattern, the F1 scores of the Glasso networks were significantly greater than those of the GAM-Glasso networks, though the magnitude of the differences in performance were minor (Figures S2, S4, and S6; Panel A). The GAM transformation also decreased or did not alter the F1 score for a small number of the SCC networks when abrupt seasonal abundance patterns were noted in the input dataset. Again, these differences in performance were small. The lower GAM-SCC network F1 scores were most prevalent when abrupt seasonal abundance patterns were coupled with low species-species covariance (Figures S3, S5, and S7; Panels C and G). There were also some statistically significant, but very small, decreases in the GAM-SCC F1 scores relative to the SCC F1 scores when 50% of the species in the input dataset had long-term increases or decreases in abundance without having seasonal abundance patterns (0%) (Figures S3, S5, and S7; Panels E-F).

In sum, there was never a substantial decrease, and there was often a substantial increase, in network predictive power when applying the GAM transformation to the data before network inference in our network simulations.

## Discussion

Ecological co-occurrence networks can be useful for capturing complex, biotic interactions when applied to high-throughput sequencing datasets of microbial communities. However, networks can yield inaccurate associations if time-series properties (i.e. seasonality, long-term trends, and autocorrelation) are prevalent in the dataset [4, 22]. The performance of the SCC and Glasso methods used in this study tended to decrease as the number of species with a seasonal abundance pattern in a dataset increased (Figure 2, Tables S2-S4). This finding indicates that time-series networks can result in a higher number of false positive and false negative associations than networks that are constructed with datasets that lack time-series features. It is likely that the false positive associations that were detected in our time-series network outputs were indirect associations that resulted from the shared periodicity of two or more species over time. Additionally, true species-species associations may have been missed by our network runs (false negatives) if seasonal and long-term signals overpowered the influence that other organisms had on species abundance patterns.

The GAM transformation carried out prior to network construction in this study improved SCC and Glasso network inference on average (Figure 2). The generally higher F1 scores that were noted following GAM transformation (Figure 2) suggest that the GAM model was able to successfully capture and remove many of the seasonal and long-term signals that were prevalent in the mock communities. Previous efforts have been made to account for indirect or time-dependent associations in network analyses. For example, the EnDED program (environmentally driven edge detection) uses environmental variables (e.g. temperature, salinity, etc.) to predict and remove indirect, environmentally driven associations in an ecological network after network inference has already been performed, provided that environmental metadata are available [46]. Time-ordered networks have also been used by researchers to account for time when conducting network analyses [22, 47]. In time-ordered networks, a node is created for each species at each time point in a dataset, thus allowing those networks to capture associations between species at specific points in time [47]. However, this approach makes no correction for seasonal or long-term trends. The GAM-based data transformation proposed here provides an effective tool that can be used to account for multiple time-series features prior to carrying out ecological network analyses, can be tailored to a wide variety of time-series datasets, and can be used in conjunction with any downstream networking method.

The SCC and Glasso network comparisons carried out in this study demonstrated that both of these methods have unique strengths and weaknesses and that method selection may depend on the research question(s) being asked. With Barabási-Albert networks, the GAM transformation improved the F1 score and the degree distribution of the SCC networks more than those of the Glasso networks (Figure 3). It is known that correlation-based networks tend to capture both direct and indirect associations in an input dataset [4, 9, 17], while Glasso networks can avoid capturing indirect associations [31]. The notably higher F1 scores and improvements in the degree distribution plots that were observed in the GAM-SCC networks relative to the SCC networks is presumably due to ability of the GAM to remove many of the indirect associations that would otherwise be detected in the SCC network analyses. While the GAM-SCC approach captured the real, Barabási-Albert degree distribution better than the GAM-Glasso approach, the GAM-Glasso approach was better able to capture the real, Erdős-Rényi degree distribution (Figure 4). Additionally, the average clustering coefficients of the GAM-Glasso networks were the most similar to the real, Barabási-Albert and Erdős-Rényi networks (Figures 3–4), suggesting that the GAM-Glasso networks were more similar to the real networks than the SCC networks in terms of network “clumpiness”.

The degree to which GAM transformation improved network prediction depended on four factors: (1) the underlying structure of the network, (2) the fraction of species exhibiting a seasonal abundance pattern, (3) the type of seasonal abundance pattern (i.e. gradual or abrupt), and the presence of long-term trends in species abundances over time. The GAM transformation did not substantively improve network inference when abrupt seasonal abundance patterns were prevalent (Figures S2-S7). It is possible that the smoothing functions used in the GAMs were unable to capture the some of the periodic spikes in species abundance that were noted in the abrupt, seasonal abundance patterns and therefore did not fully remove these abrupt signals. It is also possible that the GAM transformation inadvertently removed the influence that other species in the dataset had on the abundance pattern of a specific species and therefore decreased the number of true positive associations detected in the networks created using the GAM-transformed data (GAM-SCC, GAM-Glasso). The observation that the GAM method did not always improve model performance when seasonal abundance patterns were abrupt suggests an opportunity for future improvement. Generalized additive models are good at fitting smooth trends in the data, but other methods might be better at removing abrupt seasonal signals. In any event, decreases in the F1 scores under these specific conditions (Figures S2-S7) were marginal relative to the benefits obtained using the GAM transformation (Figure 2).

We did not explore whether our GAM removal method improved the performance of network analysis tools beyond SCC and Glasso, and it may be beneficial to expand the analysis to other approaches. Extended local Similarity Analysis (eLSA) which identifies time lagged associations [48] and Liquid Analysis (LA) which explores interactions between trios of variables [49–50] would likely be improved by removing seasonal signals. We note that both of these methods lack Glasso’s ability to exclude spurious associations [31]. Generating a method that can identify sparse networks, time-lagged associations and three-way interactions, while removing seasonal signals would be a clear future direction for a robust and flexible analysis of high-throughput data.

## Conclusion

The results of this study highlight the importance of considering temporal features when carrying out ecological network analyses with time-series data, given that time-dependent species abundance patterns may confound network predictions. The GAM-based data transformation presented here (NetGAM) provides a simple, yet effective tool that can be used to reduce the influence that time-series properties have on microbial abundance data prior to network construction. We published our method in a publicly available R package (https://github.com/sgleich/NetGAM) so that this data transformation can be used by other researchers in future time-series network analysis efforts. Accounting for seasonal abundance patterns, long-term trends, and autocorrelation in time-series datasets using our GAM method may substantially improve network inference. We recommend that future networking studies account for time-dependent species abundance patterns that may be prevalent in an input dataset in order to reduce the number of false positive and false negative associations that are detected through time-series network analyses.

## Acknowledgements

This work and SJG were supported by the Simons Foundation Grant P49802 (to DAC). JLW was supported by a postdoctoral fellowship Simons Foundation Award 653212. We thank Ben Tully and the members of the Bioinformatics Virtual Coordination Network (BVCN) for creating and contributing to the online platform that inspired this project.

