## Supplemental Figures 1-7 and Supplemental Table Legends for "NetGAM: Using generalized additive models to improve the predictive power of ecological network analyses constructed using time-series data"

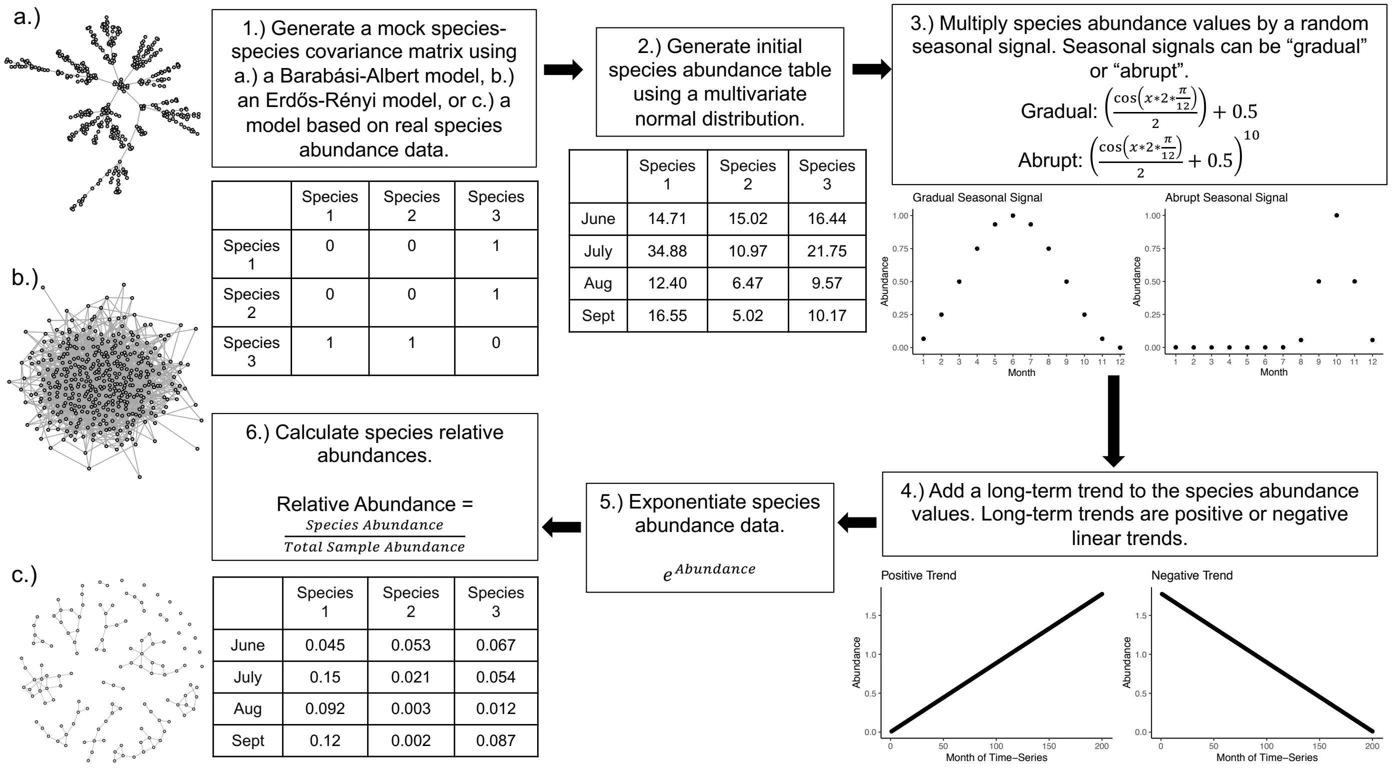


| | Species 1 | Species 2 | Species 3 |
|---|---|---|---|
| Species 1 | 0 | 0 | 1 |
| Species 2 | 0 | 0 | 1 |
| Species 3 | 1 | 1 | 0 |

| | Species 1 | Species 2 | Species 3 |
|---|---|---|---|
| June | 14.71 | 15.02 | 16.44 |
| July | 34.88 | 10.97 | 21.75 |
| Aug | 12.40 | 6.47 | 9.57 |
| Sept | 16.55 | 5.02 | 10.17 |

| | Species 1 | Species 2 | Species 3 |
|---|---|---|---|
| June | 0.045 | 0.053 | 0.067 |
| July | 0.15 | 0.021 | 0.054 |
| Aug | 0.092 | 0.003 | 0.012 |
| Sept | 0.12 | 0.002 | 0.087 |

**Figure S1:** Steps used to generate mock species abundance data with seasonal signals, long-term trends, and an underlying covariance structure. The networks labeled a-c show the 3 network structures that were tested in the GAM transformation method validation—the Barabási-Albert network (a), the Erdős-Rényi network (b), and the network predicted from the American Gut dataset (c).

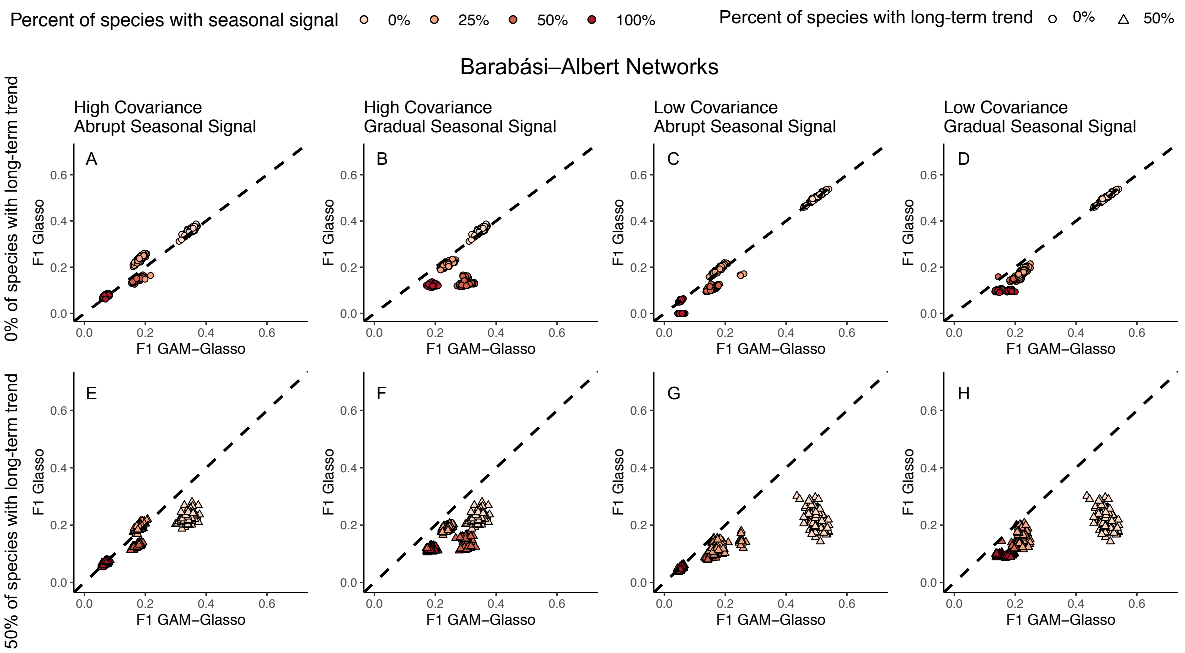


**Figure S2:** F1 score of the Glasso networking method without the GAM data transformation (F1 Glasso) as a function of the F1 score of the Glasso networking method with the GAM data transformation (F1 GAM-Glasso) for datasets in which the real network structure was that of a Barabási–Albert model. The results of 400 networks are shown in each panel. The species abundance data used in panels A-D did not have any long-term trends, while the species abundance data used in panels E-H had long-term trends added to 50% of the network species. The dashed, black lines show the 1:1 relationship. Data points below the 1:1 line depict network runs that had a higher GAM-Glasso F1 score, while data points that fall above the 1:1 line depict network runs that had a higher Glasso F1 score.

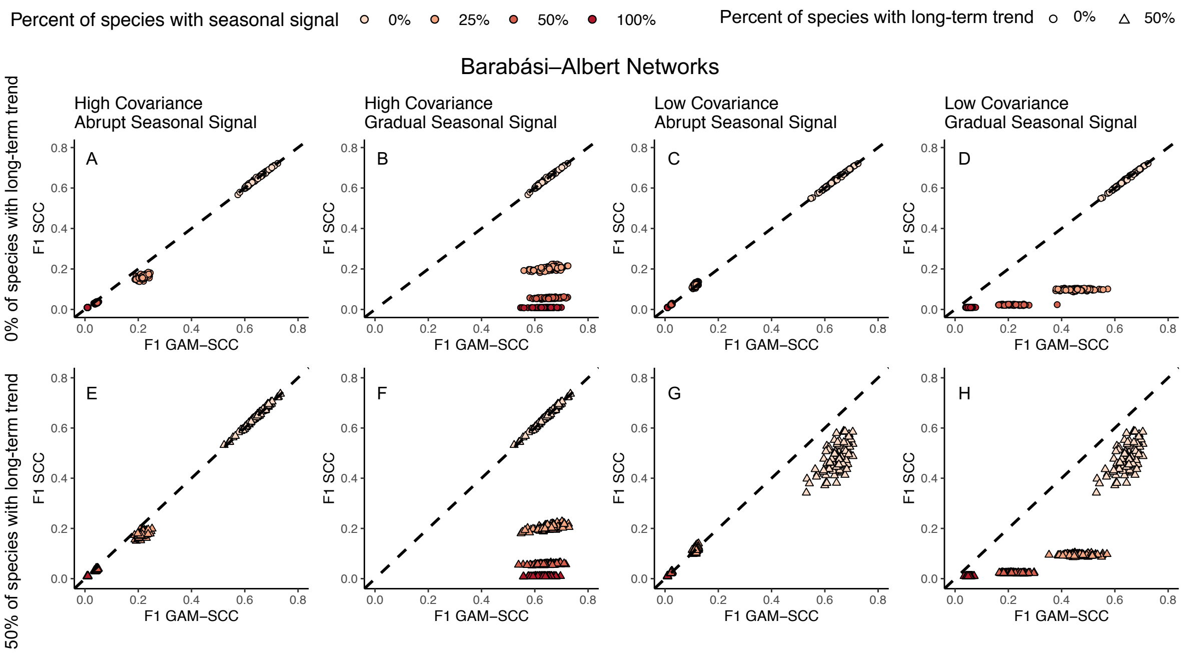


**Figure S3:** F1 score of the SCC networking method without the GAM data transformation (F1 SCC) as a function of the F1 score of the SCC networking method with the GAM data transformation (F1 GAM-SCC) for datasets in which the real network structure was that of a Barabási–Albert model. Panels and details are the same as in Figure S2.

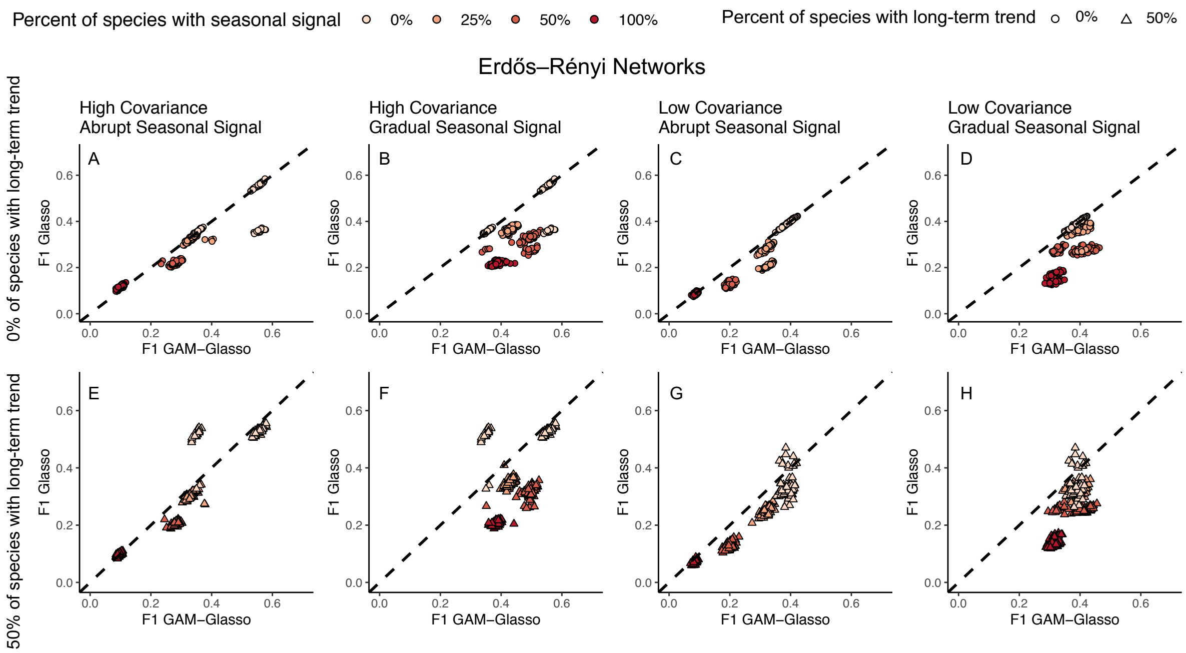


**Figure S4:** F1 score of the Glasso networking method without the GAM data transformation (F1 Glasso) as a function of the F1 score of the Glasso networking method with the GAM data transformation (F1 GAM-Glasso) for datasets in which the real network structure was that of an Erdős-Rényi model. Panels and details are the same as in Figure S2.

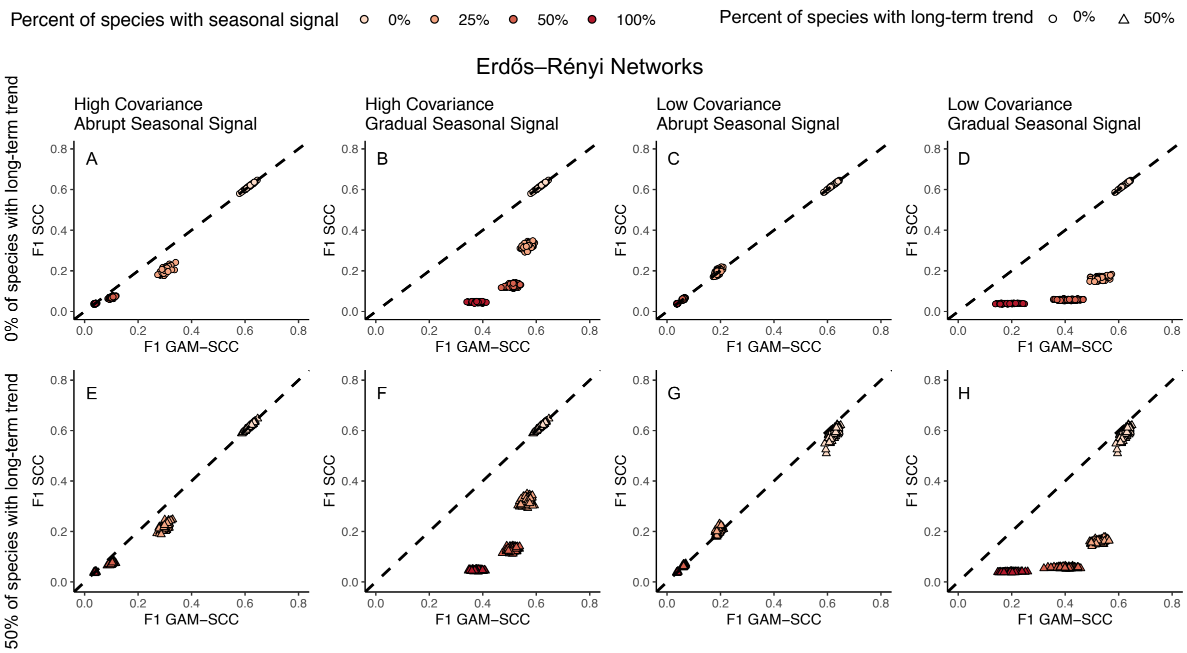


**Figure S5:** F1 score of the SCC networking method without the GAM data transformation (F1 SCC) as a function of the F1 score of the SCC networking method with the GAM data transformation (F1 GAM-SCC) for datasets in which the empirical network structure was that of an Erdős-Rényi model. Panels and details are the same as in Figure S2.

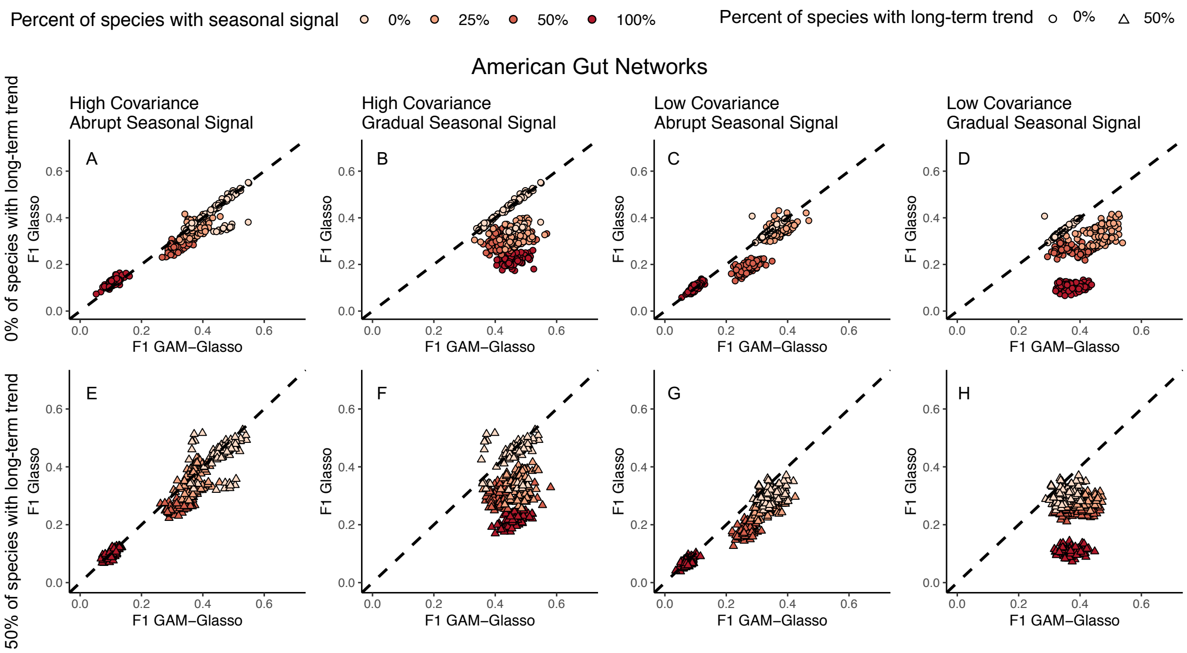


**Figure S6:** F1 score of the Glasso networking method without the GAM data transformation (F1 Glasso) as a function of the F1 score of the Glasso networking method with the GAM data transformation (F1 GAM-Glasso) for datasets in which the real network structure was estimated from the American Gut dataset. Panels and details are the same as in Figure S2.

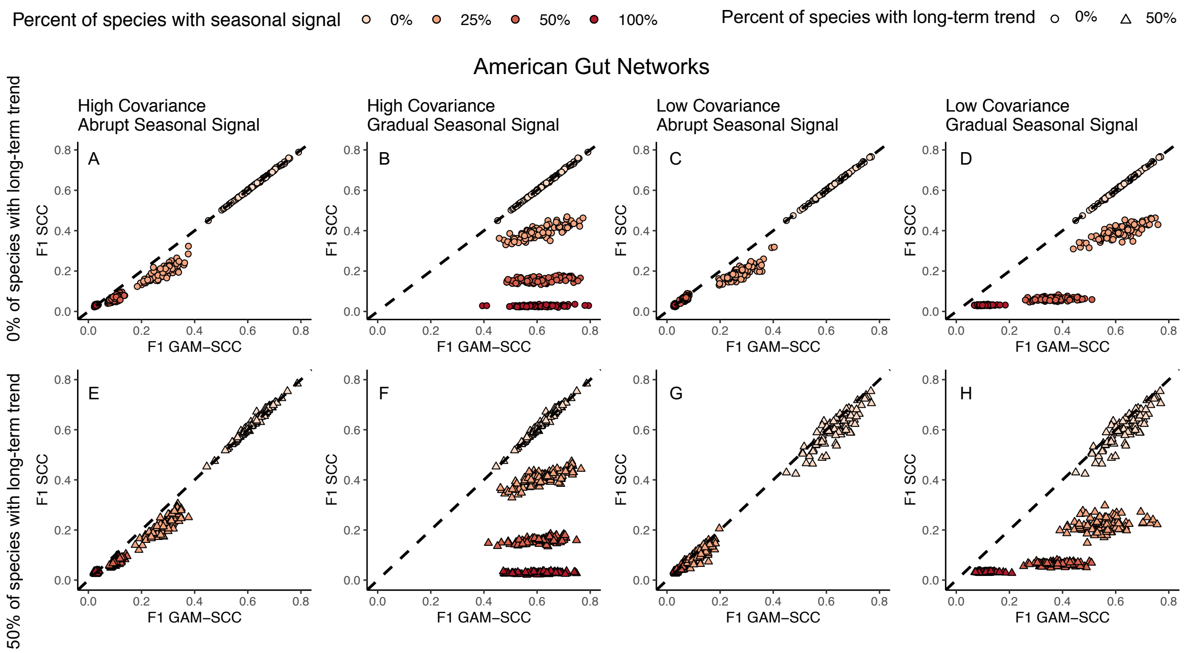


**Figure S7:** F1 score of the SCC networking method without the GAM data transformation (F1 SCC) as a function of the F1 score of the SCC networking method with the GAM data transformation (F1 GAM-SCC) for datasets in which the real network structure was estimated based on the American Gut dataset. Panels and details are the same as in Figure S2.

### Supplemental Table Legends

**Table S1:** Conditions used to generate the mock, time-series datasets that were used in this study. For each condition (row), 100 mock datasets were created. Then, the 4 (Glasso, GAM-Glasso, SCC, and GAM-SCC) network analysis approaches were used to infer species-species associations from each mock dataset. In total, 8 400 mock datasets were created, and 33 600 networks were generated (8 400 * 4).

**Table S2:** Summary statistics for the Glasso, GAM-Glasso, SCC, and GAM-SCC networks that were generated from mock datasets with a Barabási-Albert network structure.

**Table S3:** Summary statistics for the Glasso, GAM-Glasso, SCC, and GAM-SCC networks that were generated from mock datasets with an Erdős-Rényi network structure.

**Table S4:** Summary statistics for the Glasso, GAM-Glasso, SCC, and GAM-SCC networks that were generated from mock datasets with a network structure that was estimated from the American Gut dataset.
